# MicroRNA-92a negatively regulates neurofibromin 2 and inhibits its tumor suppressive function

**DOI:** 10.1101/249177

**Authors:** Krizelle Mae M. Alcantara, Reynaldo L. Garcia

## Abstract

Inactivation of the tumor suppressor Merlin leads to the development of benign nervous system tumors of neurofibromatosis type 2. Merlin deficiency is also observed in human malignancies including colorectal and lung cancers. Causes of Merlin inactivation include deleterious mutations in the encoding neurofibromin 2 gene (NF2) and aberrant Merlin proteasomal degradation. Here, we show that NF2 is also regulated by microRNAs (miRNAs) through interaction with evolutionarily conserved miRNA response elements (MREs) within its 3’-untranslated region (3‘UTR). Dual luciferase assays in HCT116 and A549 show downregulation of wild type NF2 by miR-92a via its 3’UTR but not NF2-3’UTR with mutated MRE. HCT116 cells transfected with miR-92a show significant downregulation of endogenous NF2 mRNA and protein levels, which were rescued by co-transfection of a target protector oligonucelotide specific for the miR-92a binding site within NF2-3’UTR. MiR-92a overexpression in HCT116 and A549 resulted in increased migration and proliferation, apoptosis resistance, and altered F-actin organization compared to controls. This study provides functional proof of the unappreciated role of miRNAs in NF2 regulation and tumor progression.

## INTRODUCTION

Neurofibromatosis type 2 (NF2; OMIM 101000) is a cancer predisposition syndrome which arises from inactivation of the neurofibromin 2 (*NF2*) gene with associated loss of function of its protein product Merlin (Gutmann, Hirbe, & Haipek, 2001). This autosomal dominant disorder has a birth incidence of 1 in 33,000 and a disease prevalence of 1 in 60,000 (Evans *et al*., 2010). Classical NF2 is characterized by the development of multiple benign tumors of the nervous system with hallmark symptoms of bilateral vestibular schwannomas (VS), meningiomas, schwannomas, and ependymomas (Schroeder *et al*., 2014).

Merlin regulates contact-dependent inhibition of growth via signal transduction pathways controlling cell proliferation and survival (Curto & McClatchey, 2008; Schroeder *et al*., 2014). Merlin also acts as a molecular scaffold between transmembrane receptors and the cortical actin cytoskeleton, thus regulating functions such as cell morphogenesis, adhesion, and migration (Curto & McClatchey, 2008).

Studies investigating Merlin regulation and tumorigenesis mainly focus on mutational events within the coding region of *NF2* leading to loss of functional protein expression (Morrow & Shevde, 2012). *De novo* mutations in *NF2* have been reported in 50-60% of NF2 cases (Evans, 2009b; Schroeder *et al*., 2014). Interestingly, rare somatic mutations in *NF2* have also been detected in common human malignancies not associated with neurofibromatosis type 2, including mesotheliomas, colorectal, lung, breast, hepatic, prostate, thyroid carcinomas and melanomas (Petrilli & Fernández-Valle, 2015; Schroeder *et al*., 2014; Yoo, Park, & Lee, 2012).

Despite the low prevalence of *NF2* mutations in cancer (Petrilli & Fernández-Valle, 2015), there is mounting evidence that inactivation of Merlin may be involved in cancer development and progression. Čačev *et al*. (2014) reported that *NF2* mRNA and protein expression are significantly lower in poorly differentiated colorectal carcinoma tumors compared to well differentiated tumors. In breast cancer, 75% (56 out of 75) of tumors without *NF2* gene mutations were shown to have unaltered *NF2* transcript levels but significantly low Merlin expression. This correlated with increased metastatic potential, which was reversed by rescuing Merlin expression (Morrow *et al*., 2011). These studies indicate that other molecular mechanisms may be involved in Merlin inactivation that may contribute to disease progression.

One possible mechanism is post-transcriptional regulation of the *NF2* gene by microRNAs. Endogenously expressed miRNAs have been shown to play key roles in cancer by regulating oncogenes and tumor suppressor genes through MREs within their 3’ UTR (Jannson & Lund, 2012). For Merlin, however, there is paucity of information on whether its expression and tumor suppressor function are endogenously regulated by specific miRNA species (Morrow & Shevde, 2012).

In an effort to provide more in-depth investigations on the role of miRNAs regulating *NF2*, we analyzed the 3’UTR sequence of wild-type *NF2 in silico*. Among the candidate miRNAs identified, we demonstrated that miR-92a negatively regulates *NF2* mRNA and protein expression in HCT116 colorectal cancer cells. Overexpression of miR-92a in HCT116 and A549 lung adenocarcinoma cells disrupted contact-mediated inhibition of proliferation and enhanced cell migration, proliferation, and survival. Changes in F-actin organization was also observed in miR-92a-overexpressing A549 cells. Together, our results indicate that miR-92a can impact Merlin’s tumor suppressive functions by targeting the NF2-3’UTR.

## RESULTS

### Bioinformatics analysis predicts targeting of NF2-3’UTR by hsa-miR-92a

In order to select candidate miRNAs targeting the 3’UTR of the predominant *NF2* isoform 1 mRNA (NM_000268.3), the key features of a functional miRNA:target interaction were analyzed. Context score percentile of each miRNA binding site was provided by TargetScanHuman v6.2 (http://www.targetscan.org/vert_61/). Thermodynamic stability of miRNA:target interaction was analyzed using PITA (http://genie.weizmann.ac.il/pubs/mir07/mir07_prediction.html). MirSVR scores predicting likelihood of downregulation of *NF2* by miRNAs from sequence and structure features of predicted miRNA binding sites were obtained from microRNA.org. Rank and target scores of miRNAs predicted to target *NF2*-3’UTR were analyzed by mirDB (http://mirdb.org/miRDB/).

Several miRNA families were predicted to have binding sites broadly conserved among vertebrates within the *NF2*-3’UTR sequence (Table EV1). Of these, the binding sites for hsa-miR-92a-1 (NR_029508.1) ranked highest according to aggregate preferentially conserved targeting (P_CT_) scores (Table EV1). Target-miRNA mapping showed that miR-92a has two predicted binding sites within the *NF2*-3’UTR sequence (Figure EV1), which are shared MREs with the miR-25 miRNA precursor family (hsa-miR-25, hsa-miR-32, hsa-miR-92a, hsa-miR-92b, hsa-miR-363, and hsa-miR-367). Predictive scores of the other bioinformatics tools consistently ranked miR-92a as a good potential candidate and was thus selected for this study. A summary of the bioinformatics analysis profiles of miR-92a is shown in Table 1.

**Table 1.** Summary of bioinformatics analysis profiles of hsa-miR-92a.

| miRNA | # of MREs within *NF2*-3'UTR | MRE position | $\Delta\Delta G$ (kcal/mol) | Context score percentile | P<sub>CT</sub> | MirSVR score and rank | mirDB target score and rank |
| --- | --- | --- | --- | --- | --- | --- | --- |
| hsa-miR-92a | 2 | 159-165*<br>2816-2822 | -4.43<br>-12.57 | 81<br>8 | 0.72<br><0.1 | -0.1779 (rank 5)<br>- | 52<br>(rank 146) |

\* - conserved site, broadly conserved among vertebrates

### Dual-luciferase assays suggest negative regulation of NF2 by miR-92a via its 3’UTR

To test for endogenous regulation of *NF2* via its 3’UTR, a 792-bp segment of the *NF2*-3’UTR encompassing nucleotides 26 to 818 after the stop codon was cloned into the miRNA target expression vector pmirGLO (Promega). This 3’UTR fragment includes the conserved binding site for miR-92a at position 159-165. Its shortened length limits the background effects from the pool of miRNAs potentially targeting the rest of the full-length wild-type *NF2*-3’UTR (3,798 bp) (Figure EV1). Dual luciferase assays (DLA) showed a significant decrease in normalized firefly relative luciferase units (RLUs) in HCT116 (Figure 1a) and A549 (Figure 1b) cells overexpressing wild-type *NF2*-3’UTR compared to cells transfected with vector only, suggesting that *NF2* is endogenously regulated via its 3’UTR in both cell lines.

**Figure 1.**
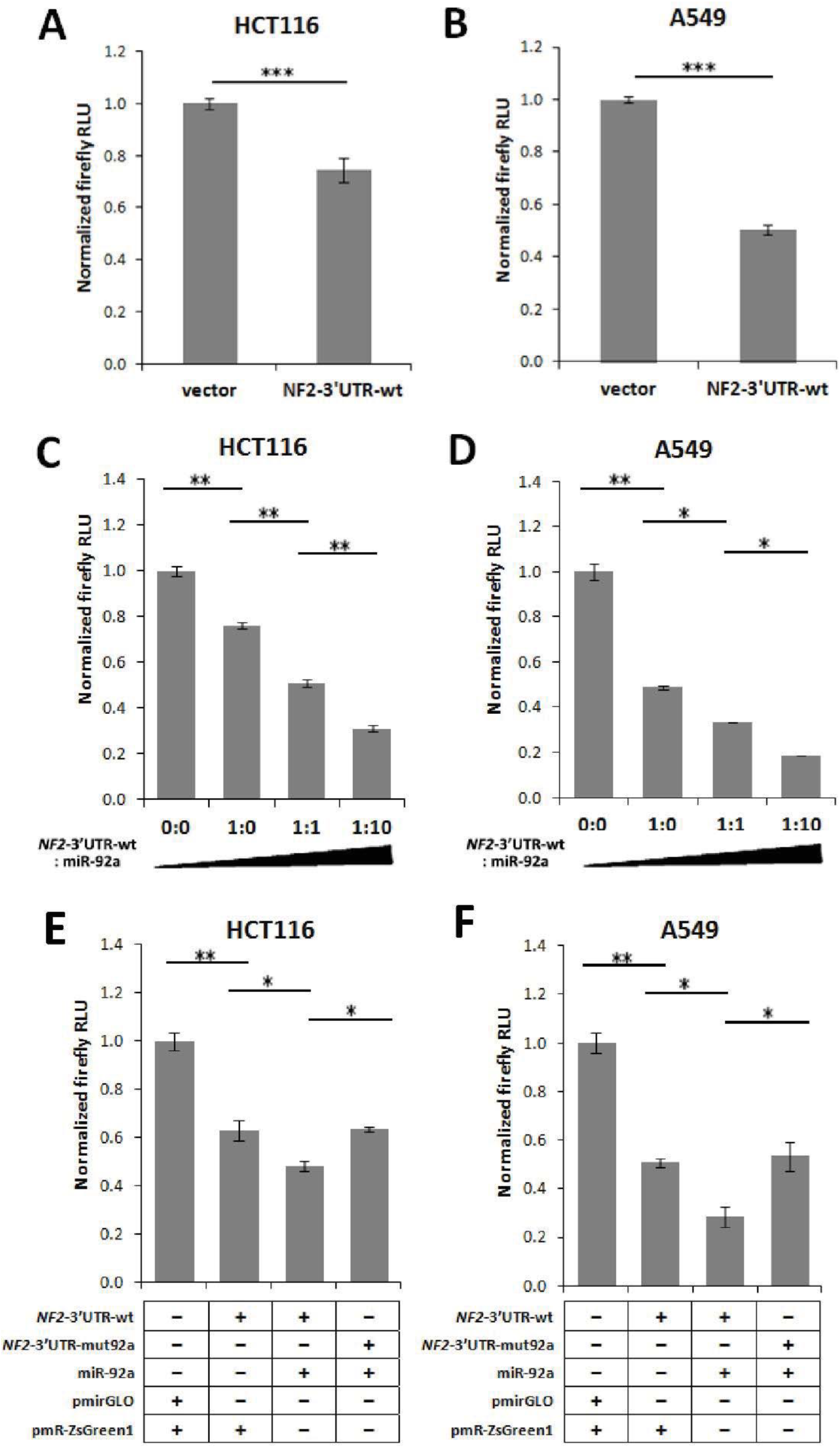
*NF2* is negatively regulated by miR-92a via its 3’UTR in HCT116 and A59 cells. *NF2*-3’UTR-wt construct and empty vector (pmirGLO) was transfected into (A) HCT116 cells and (B) A549 cells grown in 0.5% serum (n=3). Cells transfected with *NF2*-3’UTR-wt showed significantly lower luciferase reporter expression compared to vector controls. Increasing amounts of miR-92a construct were co-transfected at 0:0, 1:0, 1:1, and 1:10 *NF2*-3’UTR-wt:miR-92a plasmid ratios (ng) into (C) HCT116 cells and (D) A549 cells grown in 0.5% serum (n=3). Co-transfection of mutant *NF2*-3’UTR with miR-92a in (E) HCT116 and (F) A549 cells reversed the observed negative regulatory effect in *NF2*-3’UTR-wt setups (n=3). Data are presented as means of triplicate wells with standard deviation indicated by error bars. Firefly relative luciferase units (RLUs) were first normalized against internal control renilla luciferase RLUs before normalizing against control values. (T-test; ^*^p<0.05)

To assess whether miR-92a interacts with its conserved MRE within the *NF2*-3’UTR, co-transfection dual luciferase experiments were performed in HCT116 and A549 cells. Initial experiments for quantitation of mature miR-92a expression showed efficient processing and significant overexpression of miR-92a in transfected cells maintained in serum-depleted conditions (0.5%), but not in high serum conditions (10%) (Figure EV2). DLA results demonstrated a dose-dependent decrease in luciferase reporter expression corresponding to co-transfection of increasing amounts of miR-92a expression construct with wild-type *NF2-* 3’UTR in both HCT116 (Figure 1c) and A549 cells (Figure 1d) in serum-depleted conditions.

To confirm specificity of interaction of miR-92a with its predicted binding site within the cloned *NF2*-3’UTR segment, dual-luciferase experiments were conducted using an *NF2*-3’UTR expression construct with mutated miR-92a MRE. Analysis of the Watson-Crick base pairing of miR-92a and *NF2*-3’UTR using RNAstructure (Reuter & Mathews, 2010) showed a change in free energy (ΔΔG) of 6.1 kcal/mol between miR-92a interaction with *NF2*-3’UTR-wt (-14.2 kcal/mol) and with *NF2*-3’UTR-mut-92a (-8.1 kcal/mol) (Figure EV3). This predicts a less favorable interaction between miR-92a and the mutant *NF2*-3’UTR, leading to a derepression. Further, mutation of the shared MRE of the miR-92a family resulted to the removal of its members (miR-92a, miR-92b, miR-25, miR-32, miR-363, and miR-367) from the list of miRNAs predicted to bind to the mutant 792-bp fragment of *NF2*-3’UTR as analyzed using the MirTarget custom prediction function of miRDB (Wang, 2016). No new unintended miRNA binding sites were introduced to the mutant *NF2*-3’UTR (Table EV2).

Normalized RLUs measured from cells transiently overexpressing miR-92a co-transfected with the mutant *NF2*-3‘UTR reversed the downregulation of luciferase reporter expression in cells transfected with miR-92a and wild-type *NF2*-3’UTR, yielding luciferase readouts comparable to the empty vector controls in both HCT116 (Figure 1e) and A549 cells (Figure 1f). Overall, these results demonstrate that miR-92a negatively regulates *NF2* via its 3’UTR, in part by directly interacting with its conserved MRE within the *NF2*-3’UTR sequence.

### miR-92a negatively regulates endogenous NF2 mRNA and protein expression

MicroRNAs can regulate a gene either by target mRNA degradation and/or by translational repression. To determine whether the interaction of miR-92a with *NF2*-3’UTR affects *NF2* mRNA expression levels, semi-quantitative RT-PCR caught at the exponential phase of amplification as well as quantitative RT-PCR were performed using total cDNA generated from HCT116 cells transiently overexpressing miR-92a. Densitometric analysis of semi-quantitative RT-PCR results showed significant downregulation of *NF2* mRNA expression in HCT116 cells transfected with miR-92a compared to cells transfected with empty vector in serum-deprived conditions (Figure 2a), which corroborated with results obtained in parallel quantitative RT-PCR experiments (Figure 2b).

**Figure 2.**
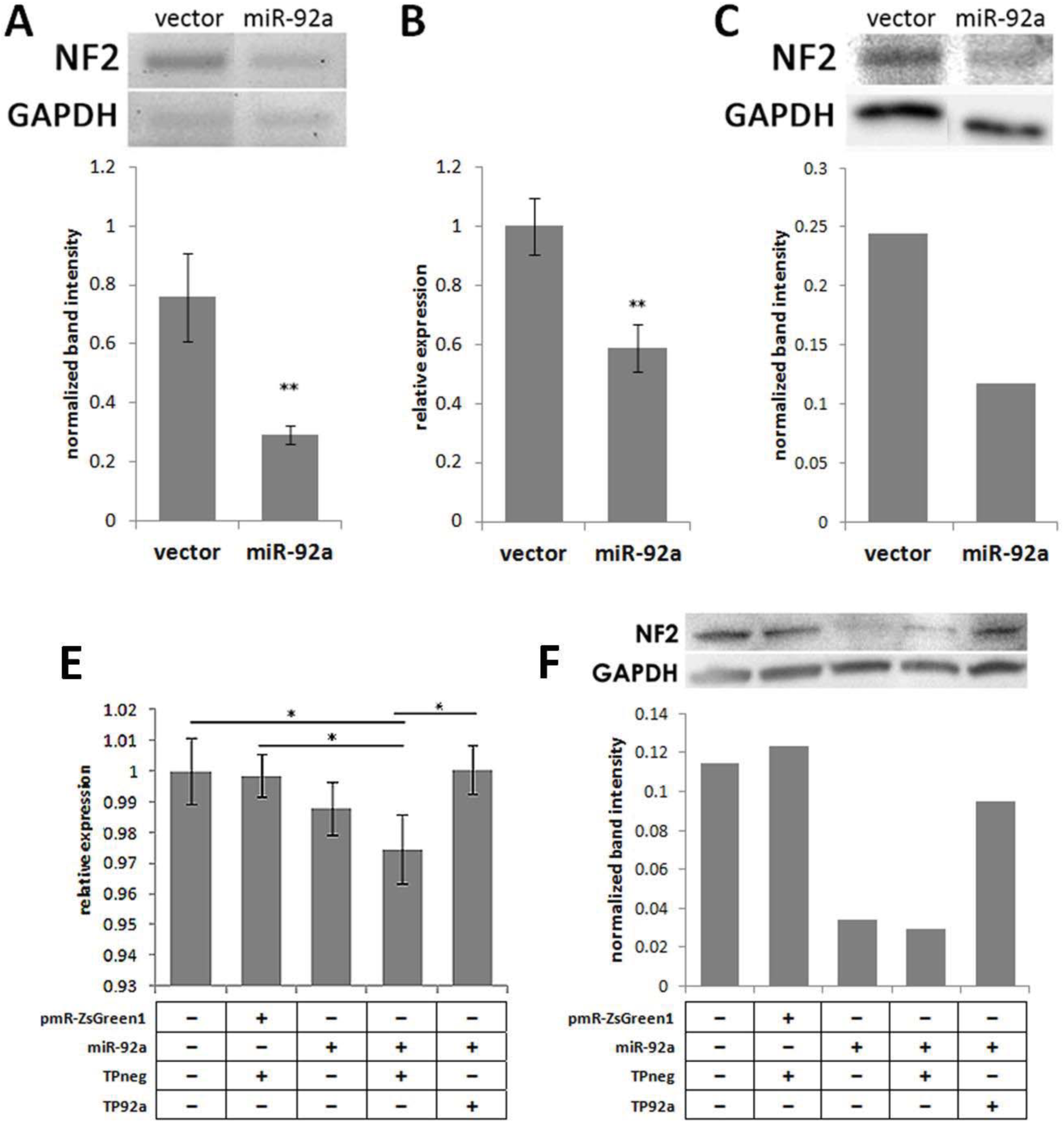
miR-92a negatively regulates *NF2* mRNA and protein levels in HCT116 cells through interaction with its conserved MRE within the *NF2*-3’UTR. (A) Semiquantitative RT-PCR gel profiles of *NF2* with GAPDH as internal control using cDNA from cells transfected with empty vector or miR-92a expression construct maintained at 0.5% FBS for 24 hrs prior to harvesting (n=3). (B) Semi-qPCR results were verified through parallel qPCR experiments (n=3). (C) Western blot of whole cell lysates from transfected HCT116 cells maintained at 0.5% FBS (n=2). HCT116 cells transfected with miR-92a have significantly downregulated *NF2* mRNA and protein expression levels. (D) qPCR and (E) Western blot was performed to probe *NF2*/Merlin expression in HCT116 cells co-transfected with pmR-ZsGreen1 and negative control target protector (TPneg), or with miR-92a expression construct and miR-92a-target protector oligonucleotide (TP-92a). *NF2* mRNA and protein expression were rescued upon co-transfection of TP-92a (n=2). Semi-qPCR and Western blot data are expressed as the average densitometric readings for *NF2*/Merlin normalized against GAPDH for all independent trials. (T-test; ^*^p<0.05)

Western blots of transfected HCT116 cells also showed markedly lower levels of Merlin protein expression in cells transiently overexpressing miR-92a versus the empty vector control as analyzed by densitometric quantification of band intensities (Figure 2c). These results demonstrate that transient overexpression of miR-92a in HCT116 cells maintained at serum-deprived conditions causes downregulation of *NF2* expression at both mRNA and protein levels, implying that miR-92a negatively regulates *NF2* through transcript degradation alone, or through translational repression as well.

Typically, miRNAs target multiple mRNAs within the cell via partial seed sequence complementarity with their respective 3’UTRs. To demonstrate that the regulatory effects of miR-92a overexpression on *NF2* mRNA and protein expression is due to direct interaction of miR-92a with *NF2*-3’UTR, a target protector oligonucleotide (TP-92a) was transfected into HCT116 cells to specifically block the conserved MRE of miR-92a within the *NF2*-3’UTR.

HCT116 cells transfected with miR-92a only or co-transfected with a negative control target protector (TPneg) display significantly downregulated *NF2* mRNA expression compared to control cells transfected with empty vector and TPneg. This negative regulatory effect was abolished upon co-transfection of miR-92a with TP-92a (Figure 2d). Similarly, co-transfection of miR-92a with TP-92a rescued the expression of Merlin which is downregulated in cells transfected with miR-92a only or co-transfected with TPneg (Figure 2e). Overall, these results demonstrate that miR-92a negatively regulates *NF2* mRNA and protein expression specifically by binding to its conserved MRE within the *NF2*-3’UTR sequence.

### Overexpression of miR-92a promotes cell proliferation and inhibits cell apoptosis

Merlin is known to function in contact-mediated inhibition of proliferation (Curto & McClatchey, 2008; Schroeder *et al*., 2014). Proliferation rates of transfected HCT116 and A549 cells were therefore tested to check whether overexpression of miR-92a can override this function of Merlin. In serum-depleted conditions (2%), HCT116 (Figure 3a) and A549 (Figure 3b) cells overexpressing miR-92a showed a significant increase in cell number between 72 hours and 96 hours post-transfection, at which point the cells are observed to reach semi-confluence (>90% growth surface area). This suggests that in space-limiting conditions, overexpression of miR-92a promotes proliferation of HCT116 and A549 cells, in part through negative regulation of *NF2* mRNA/protein expression.

**Figure 3.**
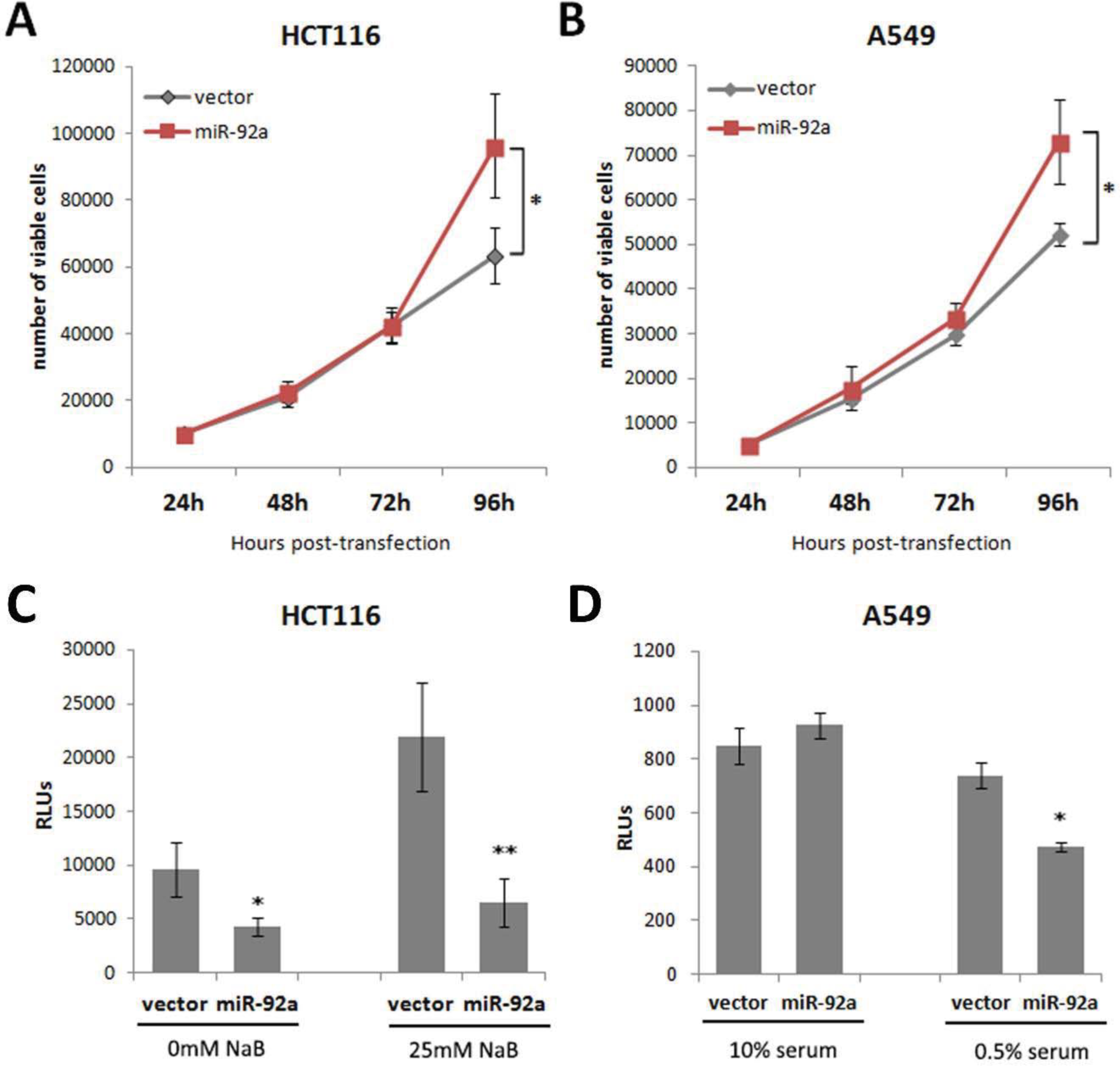
Overexpression of miR-92a enhanced cell proliferation and inhibited apoptosis induction of HCT116 and A549 cells. Representative results of triplicate experiments show HCT116 (A) and A549 (B) cells transfected with miR-92a have marked increase in proliferative capacity at 96-hours post transfection in serum-depleted conditions (2% FBS). (T-test; ^*^p<0.05). Results of triplicate experiments show the effect of miRNA overexpression on caspase 3/7 activity. (C) HCT116 cells transfected with miR-92a and fed with 2% serum showed significant decrease in caspase 3/7 activity with or without the presence of NaB. (D) A549 cells transfected with miR-92a and maintained in 0.5% serum showed significant decrease in caspase 3/7 activity. No significant effect was observed in transfected A549 cells fed with 10% serum. (T-test: ^*^p<0.05)

Merlin also exerts its tumor suppressor function by promoting apoptosis via the Hippo/SWH (Sav/Wts/Hpo) signaling pathway (Hamaratoglu *et al*., 2006). Hence, the activation of caspase-3 and caspase-7, executioner caspases of the apoptosis pathway, was measured in transfected cells to assess whether overexpression of miR-92a inhibits this function of Merlin. HCT116 cells transfected with miR-92a and maintained in low-serum conditions (2%) showed significantly lower levels of activated caspase 3/7 compared to empty vector control in both the absence or presence of the apoptosis inducer sodium butyrate (Figure 3c). Similarly, overexpression of miR-92a in A549 cells incubated in serum-deprived conditions (0.5%) displayed significantly lower levels of active caspase 3/7 compared to empty vector control (Figure 3d). Overall, these results show that miR-92a inhibits caspase 3/7 activation and promotes resistance to apoptosis in HCT116 and A549 cells, presumably in part by negative regulation of *NF2* mRNA/protein expression.

### Overexpression of miR-92a promotes cell migration

To test if overexpression of miR-92a inhibits the negative regulatory function of Merlin in cell motility, the wound healing capacity of transfected HCT116 and A549 cells was assessed. Cells overexpressing miR-92a migrated over the introduced wound gap at a significantly faster rate than cells transfected with empty vector control in both HCT116 (Figure 4a,c) and A549 cells (Figure 4b,d) in serum-depleted conditions (2%). Expression of the fluorescent ZsGreen1 marker protein was prominently observed at the migrating cell fronts, showing that positively-transfected cells were more likely to migrate into the gap. This implies that overexpression of miR-92a enhances cell motility of colorectal and lung cancer cells partly by inhibition of *NF2* mRNA/protein expression.

**Figure 4.**
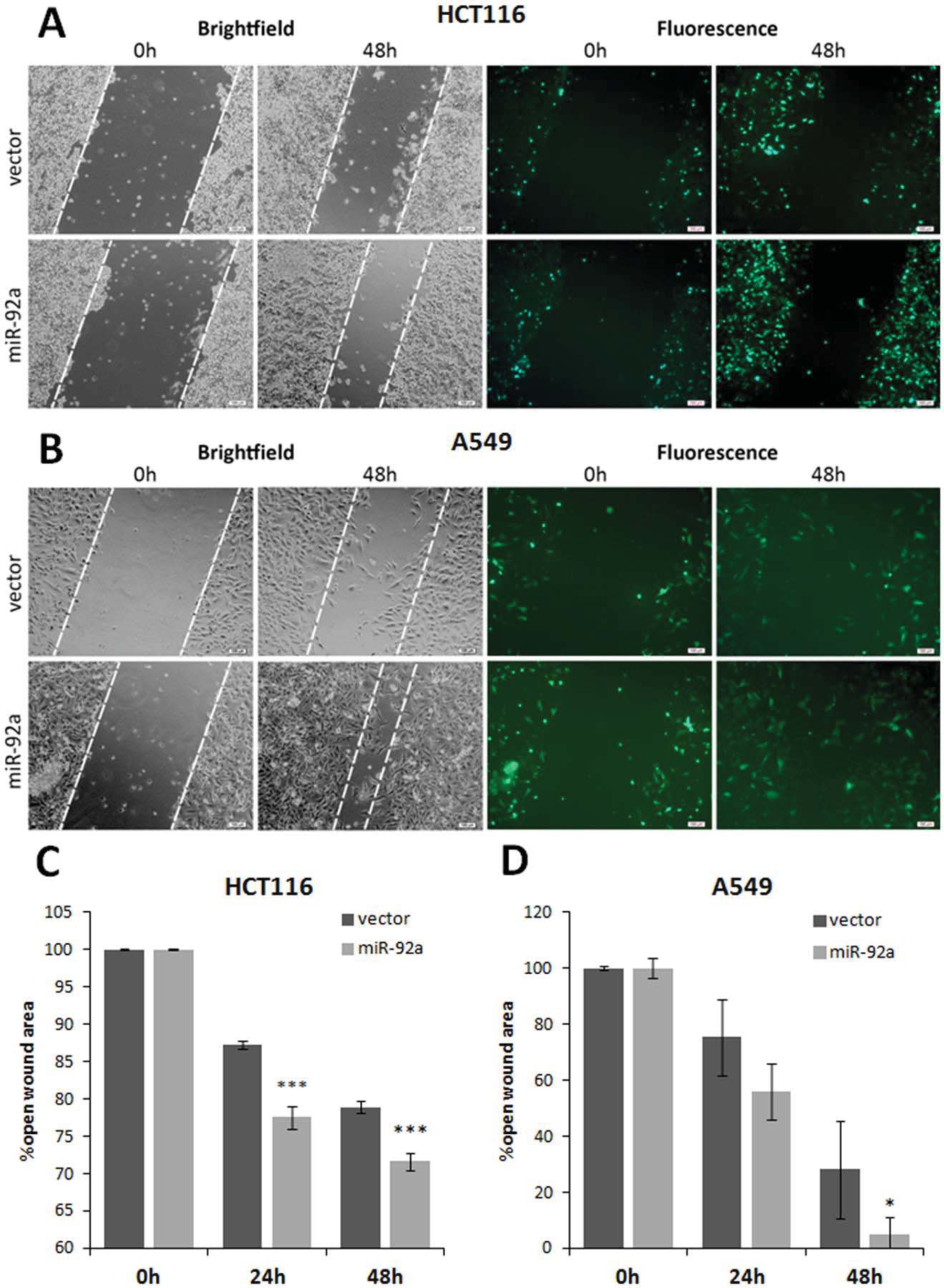
Overexpression of miR-92a increased cell motility of HCT116 and A549 cells. Wound healing assays were conducted using transfected HCT116 and A549 cells grown in 2% serum. Representative micrographs of HCT116 (A) and A549 (B) wound fields directly after scratching (0 hr) the monolayers, and 24 hrs post-scratch. Compared to the controls (i.e., vector, pmR-ZsGreen1), narrower wound gaps were observed for cells overexpressing miR-92a. Scale bars: 100 um. (C and D) Analyses of wound closure using the imaging tool Tscratch (Gebäck *et al*., 2009) shows percent open wound approximating the field view area occupied by cells at 24 hrs and 48 hrs post-scratch versus time point 0 hrs (n=3). Wound closure was significantly faster for cells overexpressing miR-92a. (T-test; ^*^p<0.05).

### Overexpression of miR-92a promotes epithelial-to-mesenchymal transition (EMT)

To explore a possible molecular mechanism behind the observed increase in migration capacity of cells overexpressing miR-92a, the expression of epithelial-to-mesenchymal transition (EMT) markers was assessed. At the transcript level, normalized band intensities from semi-quantitative RT-PCR consistently showed downregulation of *NF2* mRNA expression in cells transfected with miR-92a versus empty vector control (Figure 5a). Consequently, a decrease in the transcript expression of the epithelial marker E-cadherin was observed in miR-92a overexpressing cells, with a corresponding increase in mRNA expression of the mesenchymal marker N-cadherin (Figure 5a). Contrary to expected, a decrease in mRNA expression of vimentin was seen in miR-92a overexpressing cells (Figure 5a).

**Figure 5.**
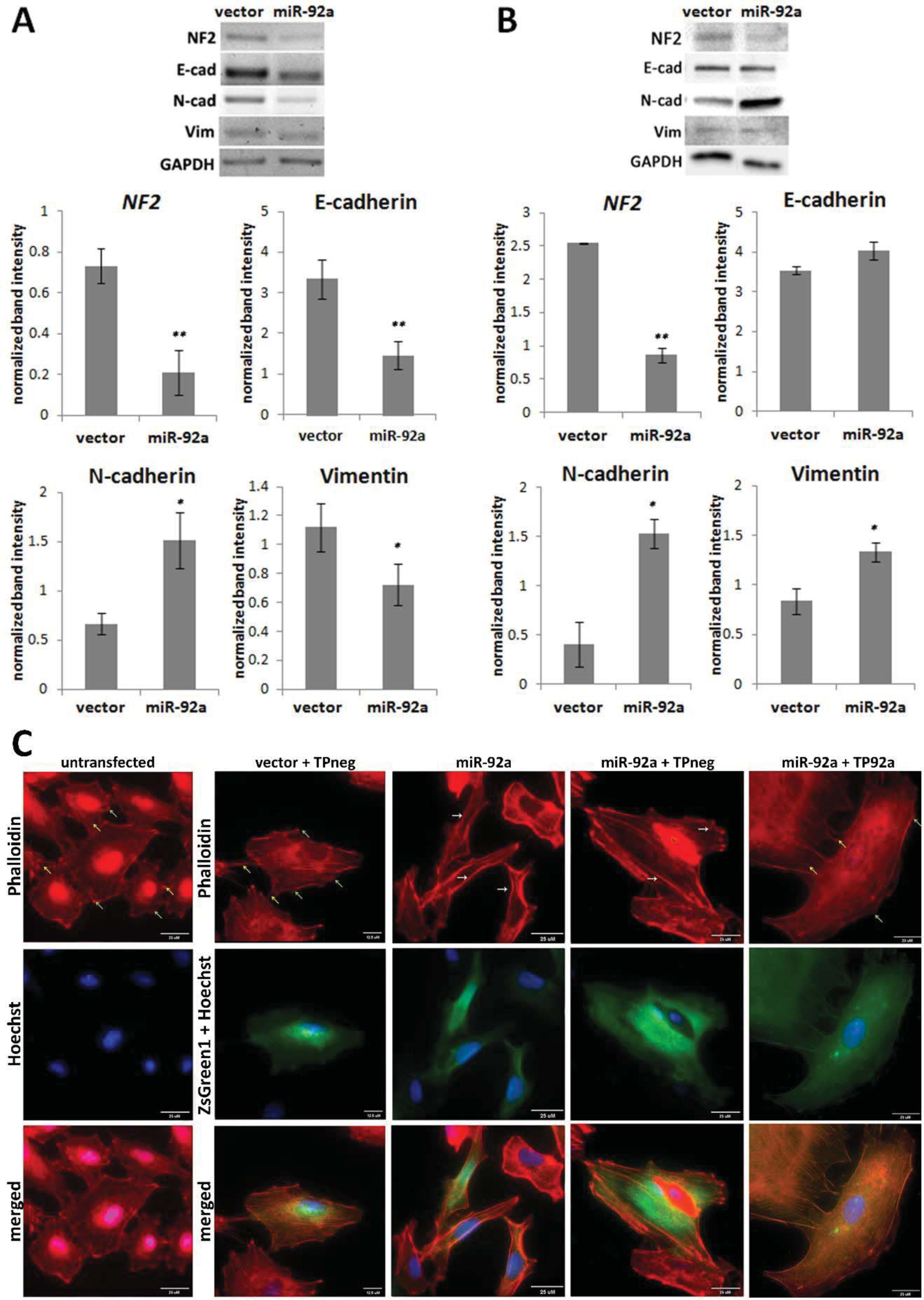
Downregulation of *NF2* by miR-92a leads to increased EMT marker expression in HCT116 cells and enhanced F-actin disorganization in A549 cells. Representative semi-quantitative RT-PCR gel profiles (A) and Western blot profiles (B) of HCT116 cells transfected with miR-92a probing mRNA expression of *NF2*, E-cadherin, N-cadherin, and vimentin. GAPDH was used as the internal control gene for normalization. Normalized band intensities show the relative expression of *NF2* and EMT marker mRNA and protein expression. Partial EMT was observed in cells overexpressing miR-92a as shown in the shift in mRNA expression between E-cadherin and N-cadherin, and in the increase in expression of mesenchymal marker proteins N-cadherin and vimentin (T-test: ^*^p<0.05). (C) Untransfected and empty vector + TPneg transfected A549 cells have notably sparse transverse actin filaments with high intercellular adhesions (yellow arrows), stable focal adhesion points (green arrows), and flat/spread-out overall morphology. In contrast, cells transfected with miR-92a, and miR-92+TPneg have dense and prominent transverse F-actin fibers (white arrows) with multiple pseudopod formation leading to a more elongated, spindle-like cell shape characteristic of motile cells. Transfection of miR-92a with a target protector for its conserved site within the *NF2*-3’UTR (TP92a) reversed the changes in morphology induced by miR-92a overexpression. (phalloidin: F-actin; Hoechst: nuclei, ZsGreen1: cells positively transfected with pmR-ZsGreen1) Scale bars: 25um.

At the protein level, normalized band intensities from Western blots showed knockdown of *NF2* protein expression in cells transfected with miR-92a compared to empty vector control (Figure 5b). No significant change in E-cadherin protein expression was observed in cells overexpressing miR-92a, although a marked increase in protein expression of both N-cadherin and vimentin was detected (Figure 5b).

Overall, these results show that knockdown of *NF2* by overexpression of miR-92a leads to an apparent shift in EMT marker expression from a more epithelial to a more mesenchymal phenotype. These results support the observed increase in migration capacities of cells overexpressing miR-92a, showing that negative regulation of *NF2* by miR-92a in HCT116 cells activates cell migration by inducing partial EMT.

### Overexpression of miR-92a alters F-actin cytoskeletal organization

Merlin is known to function as an organizer of the actin cytoskeleton primarily through the F1 and F2 subdomains within its N-terminal 4.1 ezrin-radixin-moesin (FERM) domain. Although not required *per se*, the domains also facilitate the function of Merlin in contact-inhibition of growth (Lallemand *et al*., 2009). To determine if miR-92a overexpression has deleterious effects on the cytoskeletal dynamics of epithelial carcinoma cells, the filamentous actin network of transfected A549 cells were visualized through fluorescence staining.

Untransfected A549 cells displayed a typical resting epithelial cell morphology, with an overall flat, spread-out cell shape with sparse transverse stress fibers and high intercellular cohesion. Similar F-actin organization was also observed in control cells transfected with empty pmR-ZsGreen1 vector and negative control target protector oligonucleotide (Figure 5c).

On the other hand, A549 cells transfected with miR-92a only or co-transfected with negative control target protector displayed gross morphological changes in cytoskeletal organization as evidenced by cytoplasmic shrinkage and a more spindle-shaped morphology, which are characteristics of highly motile cells (Figure 5c). Co-transfection of miR-92a with the target protector for its conserved MRE within *NF2*-3’UTR seemed to abolish the cytoskeletal changes induced by miR-92a overexpression (Figure 5). These results altogether suggest that miR-92a induces a highly dynamic actin network typical of cells with a more motile phenotype, presumably in part through negative regulation of *NF2* mRNA/protein expression.

## DISCUSSION

Deficiency in the expression levels of the tumor suppressor protein Merlin frequently occurs in multiple cancer types at a rate greater than what can be predicted by mutational analysis of the NF2 gene (Quan et al., 2015). The molecular mechanisms responsible for this are not entirely known and the consequences of Merlin downregulation in cancer progression are only recently being unraveled. In the absence of inactivating mutations in *NF2*, post-translational modifications such as Merlin phosphorylation, and epigenetic regulation such as *NF2* promoter methylation, were documented to be altered in some types of cancer (Čačev et al., 2014; Morrow et al., 2011; Quan et al., 2015). Loss of Merlin in breast, pancreatic, and colorectal cancers correlated with tumor grade associated with enhanced activation of the RAS pathway (Bakker et al., 2015; Čačev et al., 2014; Couderc et al., 2016). While post-translational and epigenetic modes of regulation could account for the disparity between the lack of mutations in *NF2* and low Merlin expression in such cases, these mechanisms have not been consistently observed across different cancer types (Morrow & Shevde, 2012). This warrants an investigation on other molecular mechanisms that may contribute to Merlin downregulation.

In this study, the potential of miRNAs to target and regulate *NF2* expression via its 3’UTR was explored. The altered expression of oncogenes, tumor suppressor genes, and cell cycle regulatory genes due to miRNA regulation are known to contribute to tumorigenesis. Here, hsa-miR-92a was identified as a novel miRNA species that regulates Merlin expression through interaction with an evolutionarily conserved MRE within the *NF2*-3’UTR. A significant decrease in luciferase expression was observed when wild-type *NF2*-3’UTR reporter construct was co-transfected with miR-92a. Mutation of this evolutionarily conserved miR-92a seed sequence abolished repression. Further, quantitative RT-PCR and Western blot analyses confirmed that miR-92a can downregulate the endogenous NF2 target by binding to this seed sequence. This was validated by target protection experiments, thus confirming the accessibility of this seed sequence within *NF2*-3' UTR to miR-92a targeting.

It should be noted that a second seed sequence of miR-92a was also identified within the *NF2*-3’UTR, which is predicted to have higher thermodynamic stability to interact with miR-92a than the conserved MRE considered for investigation in this study. However, it does not seem to significantly contribute to the negative regulation of endogenous Merlin levels, since mutation or target protection of the conserved MRE alone sufficiently abrogated the repressive effect of miR-92a overexpression on Merlin. This is consistent with the fact that this second MRE is not an evolutionarily conserved seed sequence and that it is found in the third quartile (nt positions 2816-2822 after the stop codon) of the relatively long (3,798 bp) *NF2*-3’UTR, a region less hospitable for effective targeting compared to the first MRE in the first quartile (nt positions 159-165 after the stop codon) (Grimson et al., 2007). Nevertheless, studies on this second seed sequence of miR-92a in the *NF2*-3’UTR is recommended to further establish that the miR-92a negative effect on luciferase levels is mediated primarily through its interaction with the evolutionarily conserved seed sequences.

Functional analyses demonstrated a positive role for miR-92a in promoting the metastatic properties of HCT116 colorectal and A549 lung epithelial carcinoma cells. This is in agreement with previous data showing that miR-92a has an oncogenic function in colorectal cancer (Yamada et al., 2013; Zhou et al., 2013) and lung cancer (Ren et al., 2016) cell lines and tumors. Expression levels of miR-92a in these types of cancer have been shown to correlate with their metastatic capacity (Ren et al., 2016; Zhou et al., 2013). MiR-92a expression in HCT116 and A549 is significantly higher compared to their normal cell line counterparts, FHC and BEAS-2B (Ren et al., 2016; Zhang et al., 2014). Merlin expression, on the other hand, is significantly lower in colorectal and lung cancer cells (Čačev et al., 2014; Morrow & Shevde, 2012).

In this study, exogenous overexpression of miR-92a in HCT116 and A549 cells resulted to a significant increase in proliferation and migration capacities as well as resistance to apoptosis. Given the documented role of miR-92a on NF2 regulation as described above, it would be safe to surmise that it contributes, at least partially, to the phenotypic readouts observed. Notably, the oncogenic role of miR-92a in colorectal and lung cancer cells has previously been demonstrated to be brought about partially by its negative regulation of the PTEN tumor suppressor gene (Ke et al., 2015; Ren et al., 2016) and the Kruppel-like facto4 4 (KLF4) transcription factor (Lv et al., 2016). Expression of PTEN was found to be inversely correlated with miR-92a expression in colorectal and lung cancer tissues (Ke et al., 2015; Ren et al., 2016; Zhang et al., 2014). Similar to PTEN, *NF2* is a tumor suppressor that negatively regulates the phosphatidylinositol 3-kinase/protein kinase B (PI3K/Akt) pathway by binding to phosphoinositide-3-kinase enhancer-L (PIKE-L) and thus preventing its coupling and activation of PI3K (Cooper & Giancotti, 2014; Rong et al., 2004). This suggests that the PI3K/Akt pathway is inhibited by miR-92a in colorectal and lung cancer cells, at least partially by negative regulation of PTEN and its co-target Merlin. In colorectal cancer cells, the transcription factor Kruppel-like factor 4 (KLF4) has been shown to be another co-target. Upregulation of miR-92a promoted cell proliferation and migration by direct miR-92a targeting of KLF4 (Lv et al., 2016).

The impact of miR-92a negative regulation of Merlin on its role in actin cytoskeletal organization was also assessed in this study. Cytochemical visualization of the actin cytoskeleton in A549 cells transfected with miR-92a showed cytoplasmic shrinkage and a more spindle-like morphology characteristic of highly motile cells, which is consistent with the phenotype expected if Merlin is inactivated (Sarkar et al., 2015). These changes were abolished when miR-92a was co-transfected with target protectors for its seed sequence in the NF2-3’UTR. This demonstrates the morphological transforming potential of miR-92a in part by inactivation of Merlin.

To our knowledge, this is the first *in vitro* study to functionally demonstrate the contribution of miRNAs in the regulation of Merlin’s tumor suppressive functions, in the context of colorectal cancer. An extensive literature review also revealed that apart from this report, only the work done by Perez-Garcia et al. (2015) explored the role of miRNA regulation in Merlin function, in the context of lung cancer. This study suggests that Merlin inactivation can indeed occur in cancer in the absence of deleterious mutations in the *NF2* gene. Similar studies in breast and pancreatic cancers will be instructive, since aberrant Merlin inactivation in the absence of NF2 gene mutations has been reported in these cancer types (Morrow et al., 2011; Quan et al., 2015).

## MATERIALS AND METHODS

### Cell line maintenance and transfection

HCT116 colorectal carcinoma cells and A549 lung adenocarcinoma cells were sourced from the American Type Culture Collection (ATCC). HCT116 cells were cultured in Roswell Park Memorial Institute 1640 (RPMI-1640) medium supplemented with 10% fetal bovine serum (FBS), 50 U/mL penicillin/streptomycin, and 2.0 g/L sodium bicarbonate, while A549 cells were cultured in Dulbecco’s Modified Eagle Medium:Nutrient Mixture (DMEM) supplemented with 10% FBS, 50 U/mL penicillin/streptomycin, and 3.7 g/L sodium bicarbonate.

All transfection experiments were performed using Lipofectamine^®^ 2000 (Invitrogen, MA, USA) as per manufacturer’s instructions. The amount of plasmid and volume of Lipofectamine 2000 was optimized to achieve 70-80% transfection efficiency for all functional and molecular characterization experiments (2000ng:4uL per well in 12-well plates, and 220 ng:0.4uL per well in 96-well plates).

### Antibodies and Plasmid constructs

The rabbit polyclonal anti-*NF2* (PA5-35316) and mouse monoclonal anti-N-cadherin (MA5-15633) antibodies were from Invitrogen. The rabbit polyclonal anti-E-cadherin (07-697) and mouse monoclonal anti-GAPDH (CB1001) antibodies were from EMD Millipore. The rabbit polyclonal anti-vimentin (SAB4503083) antibody was from Sigma.

The 3’UTR of human *NF2* isoform I (NM_000268.3) and the pre-miR-92a gene (NR_029508.1) were amplified by PCR from human genomic DNA using primers listed in Table S3. The *NF2*-3’UTR fragment encompasses nucleotide positions 2257 to 3048 of the variant I mRNA sequence and contains the predicted microRNA response element (MRE) hsa-miR-92a-1 at positions 159-165 of the 3’UTR. The wild-type *NF2*-3’UTR fragment was cloned into the pmirGLO dual-luciferase miRNA target expression vector (Promega), downstream of the firefly luciferase gene. A mutant version of the *NF2*-3’UTR was generated by site-directed mutagenesis of the wild-type *NF2*-3’UTR in pmirGLO, in which a CA>GT substitution within the miR-92a-1 seed sequence binding site (5’-GTG<u>CA</u>AT-3’) was introduced through overlap-extension PCR using the primers listed in Table S3.

The miR-92a precursor primers were designed to target the genomic DNA template within a range of 100-300 bp of the regions flanking the pre-miRNA sequences. The pre-miR-92a amplicon was cloned into the mammalian microRNA expression vector pmR-ZsGreen1 (Clontech).

### Quantigene evaluation of miRNA Expression in HCT116 and A549 cells

For direct quantification of mature miRNA targets in control and transfected HCT116 and A549 cells, QuantiGene miRNA Assay (Affymetrix) was used. Quantigene Singleplex miRNA probes were obtained from the pre-designed probes offered by eBioscience (Affymetrix, CA, USA). The custom probe for miR-92a (5’-UAUUGCACUUGUCCCGGCCUGU -3’) has previously been functionally validated for sensitivity and specificity for mature target miRNAs in human samples, and also evaluated for cross-reactivity towards closely-related miRNA family members.

Cultured HCT116 and A549 cells were lysed to release and stabilize miRNAs, followed by overnight (16-20 hrs) hybridization in a 96-well plate with target-specific capture probes, capture extenders, and label extenders. Pre-amplifier molecules were added and allowed to hybridize to the miR-92a-specific probe sets for signal amplification. Amplifier molecules were hybridized with pre-amplifiers, which were then probed with oligonucleotides conjugated with alkaline phosphatase (AP). A chemilumigenic AP substrate was added and the luminescent signal generated was measured using a plate-reading luminometer (FLUOstar Omega Microplate Reader, BMG LABTECH).

Triplicates of cell lysates per set-up of untransfected cells (maintained in 10% and 0.5% serum) and miR-92a-transfected cells were subjected to the assay. Average signal-background (S-B) of samples were interpolated in a standard curve generated using serial dilutions of control miRNAs to calculated target miRNA expression in each experimental setup. Average mature miRNA expression was calculated per setup.

### Dual-luciferase assay

HCT116 cells were seeded at 10,000 cells/well of a 96-well plate and transfected after 24 hrs. For the endogenous dual-luciferase assays, cells were transfected with 200 ng of empty pmirGLO vector (Promega) or pmirGLO-*NF2*-3’UTR-wt. For the miRNA co-transfection dual-luciferase assays, cells were co-transfected with 20 ng of empty pmirGLO vector, pmirGLO-*NF2*-3’UTR-wt, or pmirGLO-*NF2*-3’UTR-mut92a together with 200 ng of empty pmR-ZsGreen1 vector (Clontech) or pmR-ZsGreen-miR-92a construct. The construct preparations were complexed with Lipofectamine 2000 for transfection of cells in triplicate wells. Luciferase activity was determined 48 hrs post-transfection using the Dual-Luciferase^®^ Reporter Assay System (Promega). Data is expressed in mean values of normalized of Firefly relative luciferase units (RLUs) per setup. Raw firefly RLUs were first normalized against internal control renilla luciferase RLUs per well before normalizing against values obtained for the empty vector control setup.

### Target protector experiments

HCT116 cells were co-transfected with pmR-ZsGreen1-miR-92a and a target protector oligonucleotide specific to the conserved MRE of hsa-miR-92a within the 3’UTR of the *NF2* gene. The oligo was designed using Qiagen’s miRNA Target Protector design tool (https://www.qiagen.com/ph/shop/genes-and-pathways/custom-products/custom-assay-products/custom-mirna-products/#target-protector) using the RefSeq ID of *NF2* transcript variant 1 (NM_000268) as reference template. An effective concentration of 100nM miR-92a target protector co-transfected with ZsG-miR-92a (final concentration 2ng/uL) was determined through preliminary transfection optimization experiments. Efficiency of miRNA inhibition by the target protector was measured using quantitative RT-PCR or Western blot of lysates from transfected cells versus control setups.

### Quantitative reverse transcription PCR (qRT-PCR)

HCT116 cells were seeded at 300,000 cells/well of a 12-well plate and transfected after 24 hrs with 2000 ng of empty pmR-ZsGreen1 or pmR-ZsGreen1-miR92a construct using Lipofectamine 2000. Cells were harvested 48 hrs post-transfection and total RNA was extracted using TRIzol^®^ reagent (Invitrogen). First-strand cDNA was generated from 2000 ng total RNA per setup using M-MLV reverse transcriptase (Promega) and oligo-dT primers following manufacturer protocol. The first strand cDNA samples were then used as template for semi-quantitative RT-PCR and quantitative RT-PCR with SYBR^®^ Select Master Mix (Invitrogen) using the relative quantitation method. For semi-qRT-PCR, exponential phases of each primer pair were determined via cycle optimization PCR, followed by a second round of PCR using the optimized cycling parameters. Densitometric analysis of the digitized band intensities on the agarose gels were performed using GelQuant.NET (v1.8.2). Quantified *NF2* gene expression in triplicate experiments were normalized against GAPDH gene expression. For qRT-PCR, total *NF2* mRNA was normalized against GAPDH mRNA measured per sample. Mean values of triplicates per setup were obtained, and fold-change of *NF2* expression relative to empty vector control was calculated. Primers used for qRT-PCR are summarized in Table S3.

### Western blotting

HCT116 cells were seeded and transfected in 12-well plates as described for qRT-PCR. Cells were harvested 48 hrs post-transfection and total protein was extracted using radioimmunoprecipitation (RIPA) lysis buffer (150 mM NaCl, 0.5% sodium deoxycholate, 0.1% sodium dodecyl sulphate (SDS), 50 mM Tris pH 8.0) supplemented with protease inhibitors (1 mM phenylmethylsulfonyl fluoride (PMSF), 5 mM EDTA, and 10 μM E64 (Roche)). Lysates were clarified at 10,000g for 20 min at 4°C. Total protein concentration was measured using the bicinchoninic acid (BCA) assay. Protein extracts were snap-frozen in liquid nitrogen and stored at -80°C until use. For polyacrylamide gel electrophoresis, 30 ug of total protein per setup was loaded into Any kD^™^ Mini-PROTEAN^®^ TGX Stain-Free^™^ Protein Gels (Bio-Rad). Membranes were probed overnight with the primary antibodies described above. Upon incubation with the appropriate secondary antibody conjugated with horseradish peroxidase (HRP), bands were developed with enhanced chemiluminescence (ECL) substrate and imaged using the ChemiDoc Touch Imaging System (Bio-Rad). Densitometric analysis of digitized band intensities were performed using GelQuant.NET (v1.8.2). Quantified *NF2* gene expression were normalized against GAPDH or total protein expression.

### Wound healing assay

HCT116 cells were seeded at 300,000 cells/well and A549 cells were seeded at 200,000 cells/well of a 12-well plate. Cells were transfected 24 hrs after seeding with 2000 ng of empty pmR-ZsGreen1 or pmR-ZsGreen1-miR92a construct with Lipofectamine 2000. Upon reaching >90% confluence post-transfection, a thin artificial wound was introduced to the cell monolayer using a sterile pipette tip. Cells were washed once with 1X phosphate buffered saline (PBS), and maintained in the appropriate media supplemented with 10% or 2% fetal bovine serum (FBS). Wound closure was monitored every 12 hrs by capturing three fields of view per setup at 40X magnification using an Olympus Ix71 inverted fluorescent microscope. Percent open wound area was analyzed using the TScratch software (Gebäck et al., 2009) and reported as the mean % open wound area per setup relative to % open wound area upon initial scratching (t=0h).

### Cell proliferation

HCT116 cells were seeded at 10,000 cells/well while A549 cells were seeded at 5,000 cells/well of a 96-well plate. Cells were transfected 24 hrs after seeding with 200 ng of empty pmR-ZsGreen1 or pmR-ZsGreen1-miR-92a construct with Lipofectamine 2000 and maintained in the appropriate media supplemented with 10% or 2% FBS. The number of metabolically active cells per setup was measured at 48 hours, 72 hours, and 96 hours posttransfection upon addition of 10uL of CellTiter 96^®^ Aqueous One Cell Proliferation Assay Reagent (Promega) per well until color development. Absorbance values at 460nm of each set-up were measured with a colorimetric plate reader (FLUOstar Omega Microplate Reader, BMG LABTECH). Cell counts were calculated from a standard curve (number of cells vs A_460_) generated using serial dilutions of an untransfected cell suspension. Mean cell counts were calculated per setup for each timepoint and statistically analyzed.

### Caspase 3/7 assay

Cells were seeded and transfected in triplicate in 96-well plates as described above. HCT116 cells were then incubated in RPMI-1640 media supplemented with 10% or 2% FBS only, or with 2.5 mM sodium butyrate for induction of apoptosis. A549 cells were incubated in DMEM media supplemented with 10% FBS or without FBS. Caspase-Glo^®^ 3/7 Assay reagent (Promega) was added to each well 24 hrs post-induction. Plates were incubated at ambient temperature for 2 hrs before luminescence per well was measured with a plate reader (FLUOstar Omega Microplate Reader). Mean luminescence readings per setup were calculated per setup for each timepoint and statistically analyzed.

### Actin cytoskeleton staining

HCT116 and A549 cells were seeded into an 8-well chamber slide (Ibidi) and transfected after 24 hours. Forty-eight hours post-transfection, cells were fixed 100% methanol for 20 minutes at -20°C. PBS (1X) was used to wash cells in between the fixing, permeabilization, blocking, staining, and mounting steps. Cells were permeabilized using 0.1% Triton X-100 in 1X PBS for 15 minutes, and blocked with 1% BSA in 1X PBS for 20 minutes. For actin staining, cells were incubated in a 1:100 dilution of tetramethylrhodamine (TRITC)-conjugated phalloidin (Invitrogen). Nuclei were counterstained with Hoechst 33258 (1 μg/μL). Stained cells were mounted in 70% glycerol, covered with a glass cover slip and sealed. Stained cells were visualized under an inverted fluorescence microscope (Olympus IX83), using red fluorescent filter (λex/λem: 490/525 nm) to view stained filamentous actin structures, blue fluorescent filter (λex/λem: 355/465 nm) to view nuclei, and green fluorescent filter (λex/λem: 490/525 nm) to view cells transfected with pmR-ZsGreen1 constructs. Images were analyzed for patterns of actin arrangement in transfected compared to control or untransfected cells.

### Statistical analysis

Statistical analysis of data was performed using unpaired two-tailed T-test to measure differences between two set-ups. ANOVA with post-hoc Tukey’s honest significant difference (HSD) was used to test significant differences between multiple set-ups. Data from all quantitative experiments are presented as mean ± standard deviation (SD). In all tests, significance value was defined as ^*^p<0.05; ^**^p<0.01; ^***^p<0.001.

## Acknowledgments

This work was supported by grants from the University of the Philippines System (OVPAA-EIDR Code 06-008) and the Philippine Council for Health Research and Development (Grant Code FP150025).

## Author contributions

Conceived project: RLG. Performed experiments: KMMA. Data analyses and organization: KMMA and RLG. Wrote and edited manuscript: KMMA and RLG.

## EXPANDED VIEW FIGURE LEGENDS

**Figure EV1.**
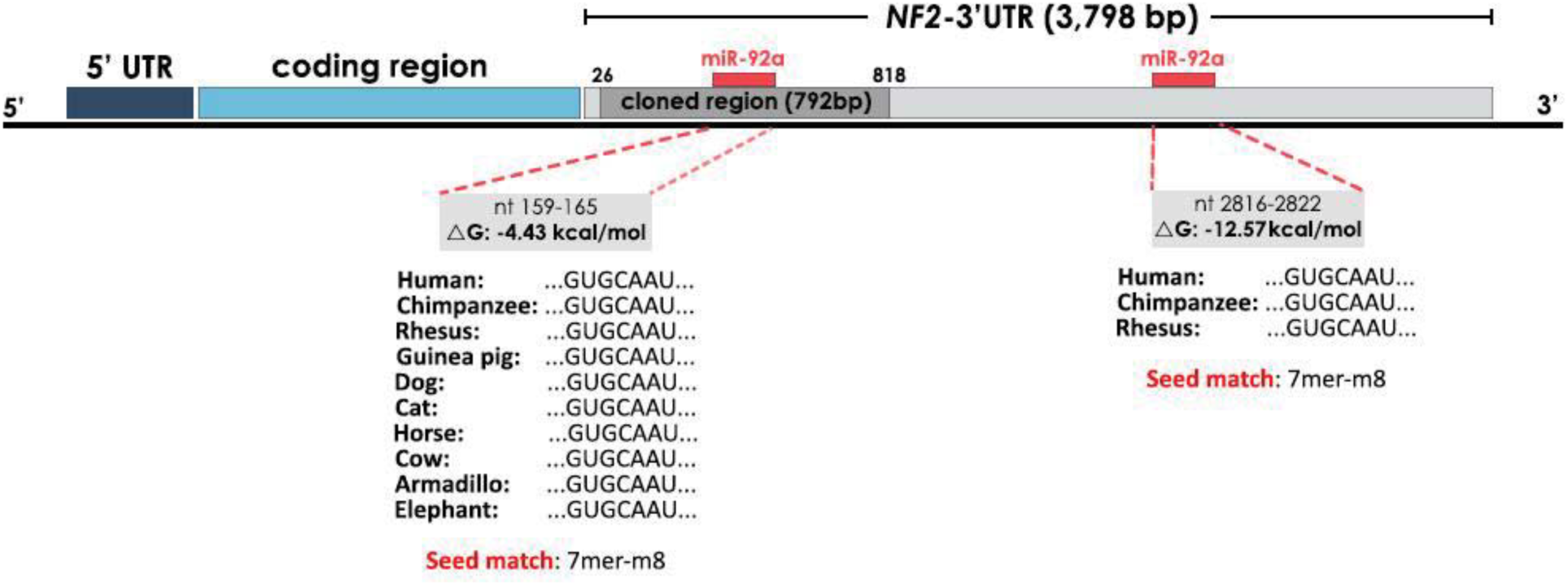
The 3'UTR of *NF2* isoform I contains evolutionarily conserved binding sites for hsa-miR-92a. Schematic representation of the human *NF2* isoform I mRNA showing the full and cloned 3’UTR region and relative positions of the predicted microRNA response elements (MREs) of hsa-miR-92a-1 at nt 159-165 and nt 2816-2822. The phylogenetic conservation and sequence complementarity (seed match) of each seed sequence is shown.

**Figure EV2.**
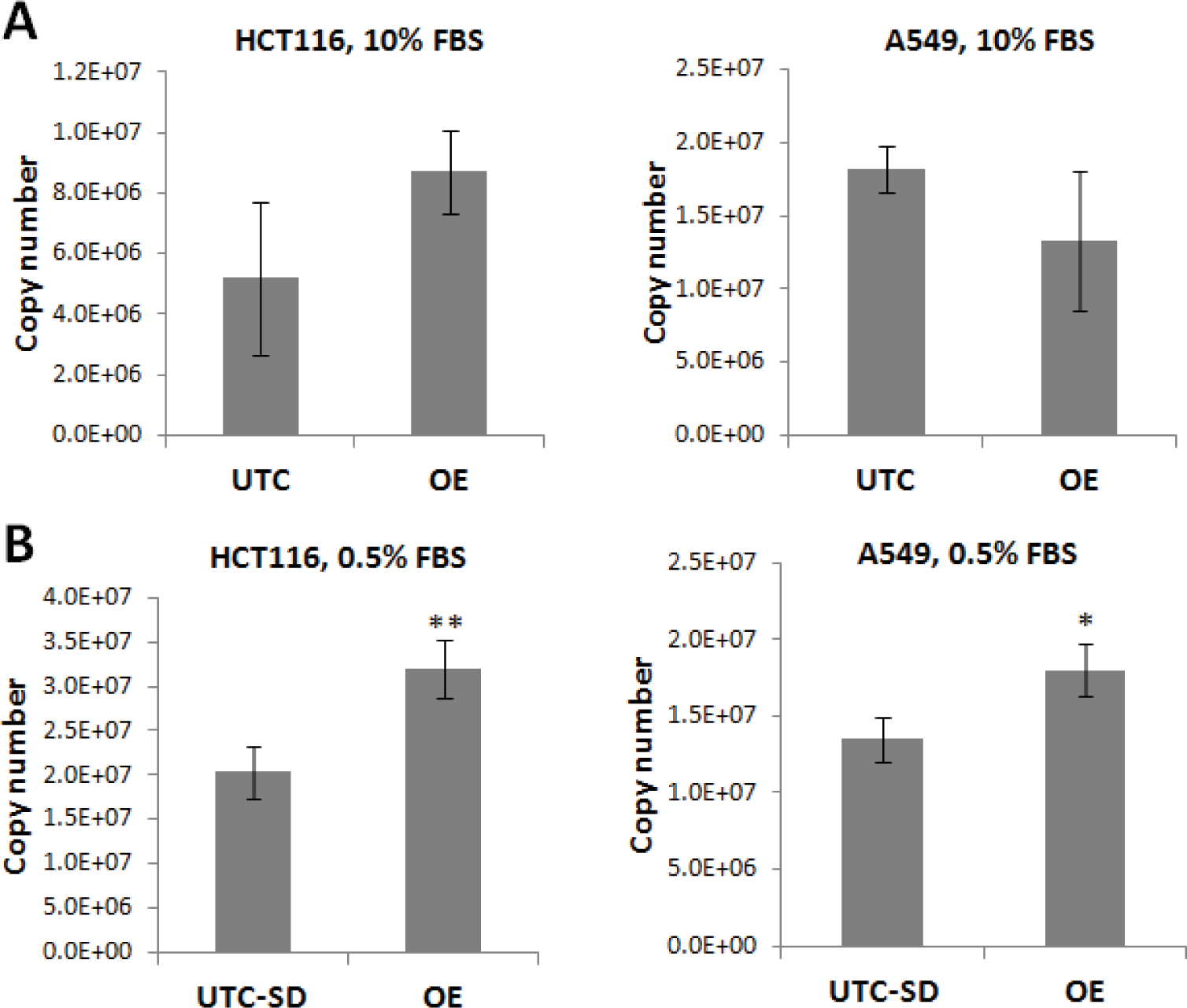
Expression levels of miR-92a was confirmed by QuantiGene miRNA assay in untransfected and transfected HCT116 and A549 cells. Mature miR-92a levels detected in untransfected and transfected HCT116 and A549 cells at 10% or 0.5% FBS. (T-test; ^*^p<0.05)

**Figure EV3.**
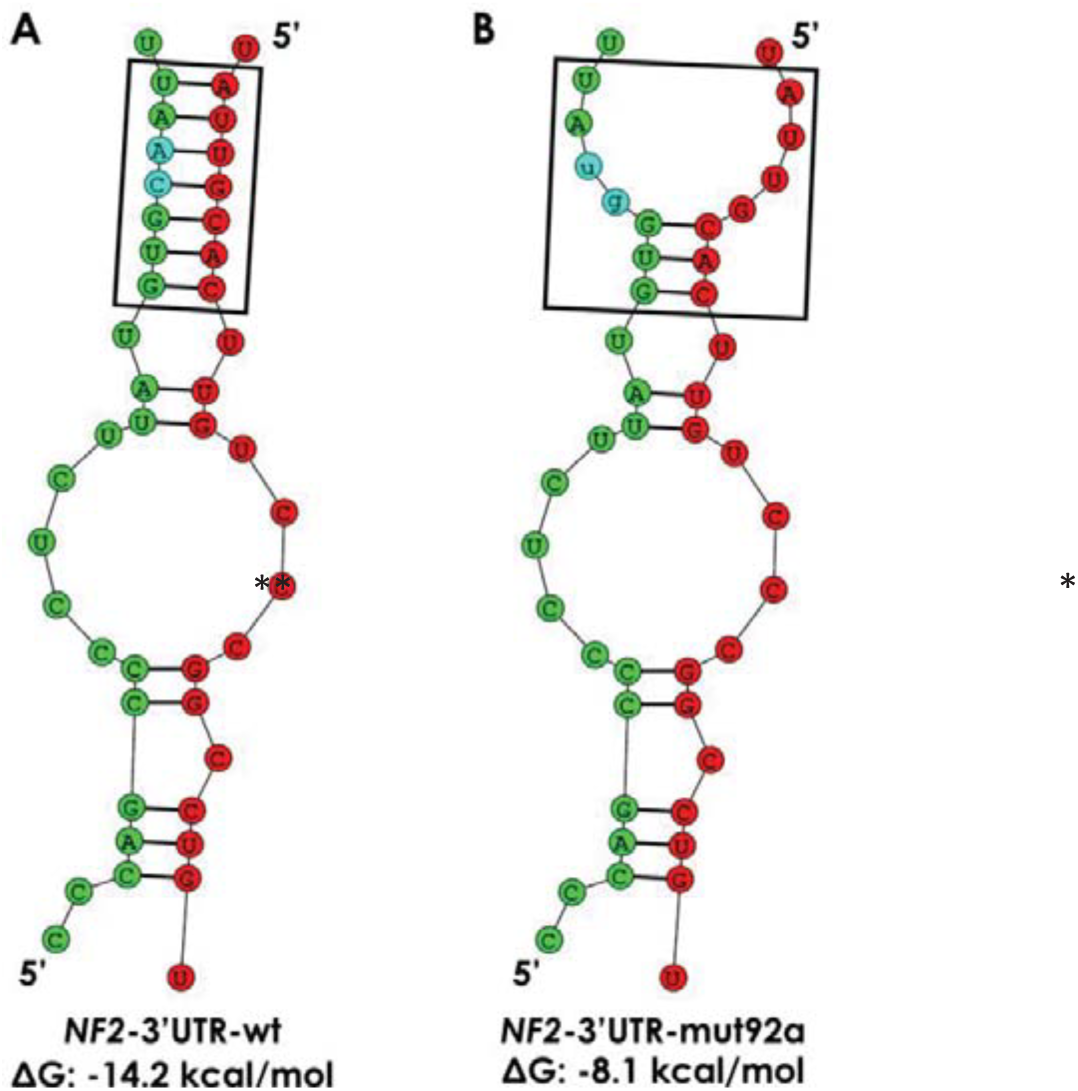
Secondary structure analysis shows the destruction of the RNA-RNA interaction between miR-92a and the predicted miR-92a MRE within the *NF2*-3’UTR upon SDM. mRNA (green)-miRNA (red) duplexes and free energy changes were predicted using RNAstructure tool set at default parameters. The 2-nt substitutions are highlighted in blue. Analysis shows significant changes in the secondary structures of the predicted interaction site (boxed region) of the miR-92a seed sequence and the *NF2*-3’UTR miR-92a MRE as well as in the free energy of interaction between *NF2*-3’UTR and miR-92a in wild-type (A; -14.2 kcal/mol) vs mutant (B; -8.1 kcal/mol) *NF2*-3’UTR.

## EXPANDED VIEW TABLE LEGENDS

**Table EV1.**
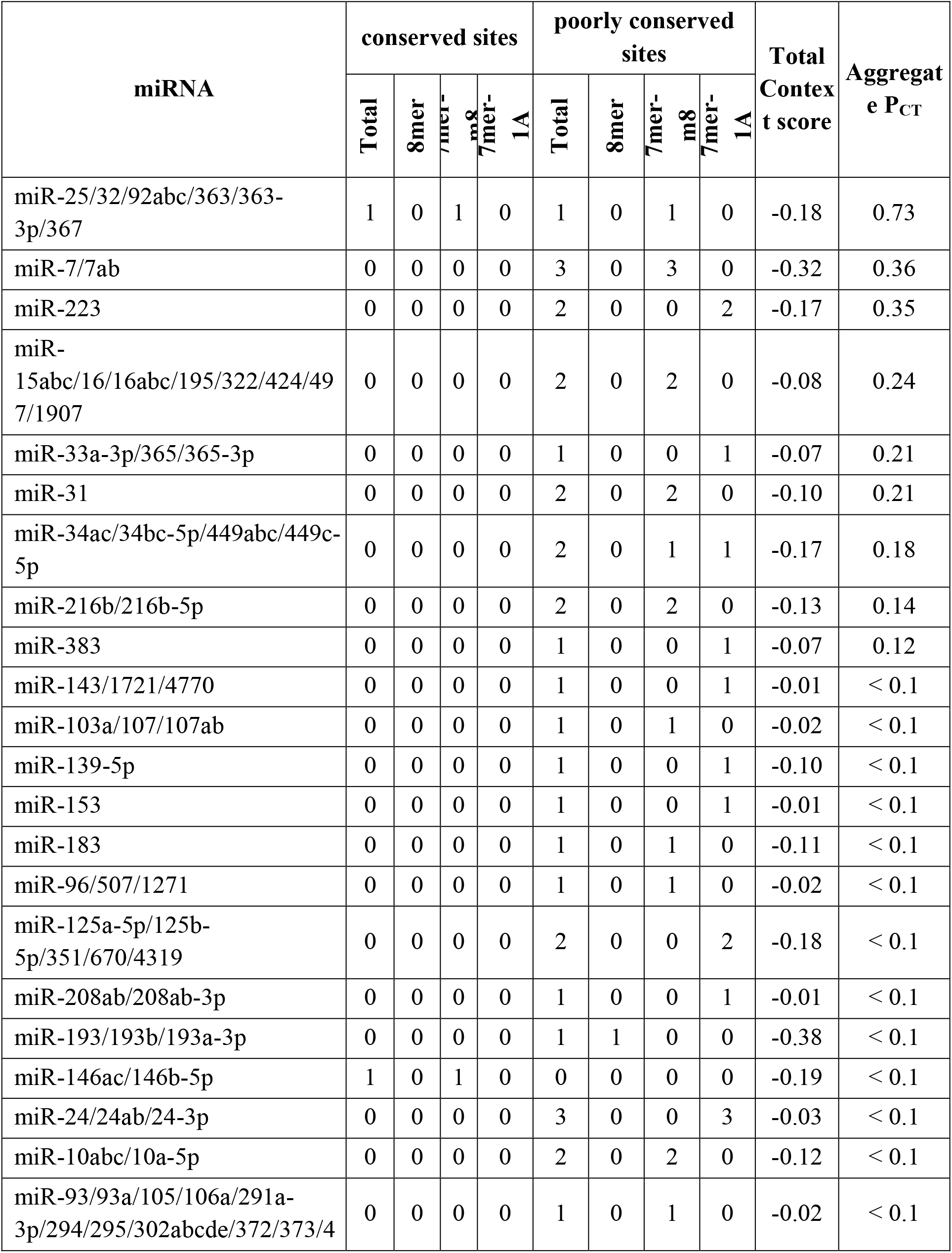
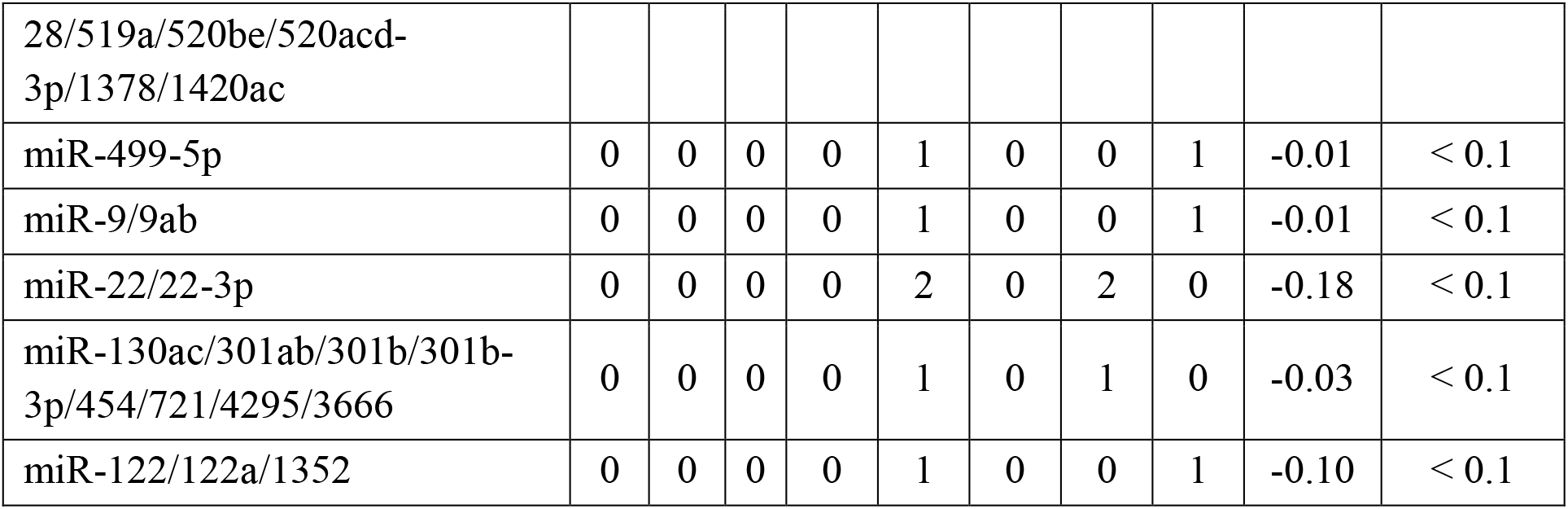
List of miRNA families targeting Human *NF2* (NM_181833) 3' UTR broadly conserved among vertebrates.

**Table EV2.**
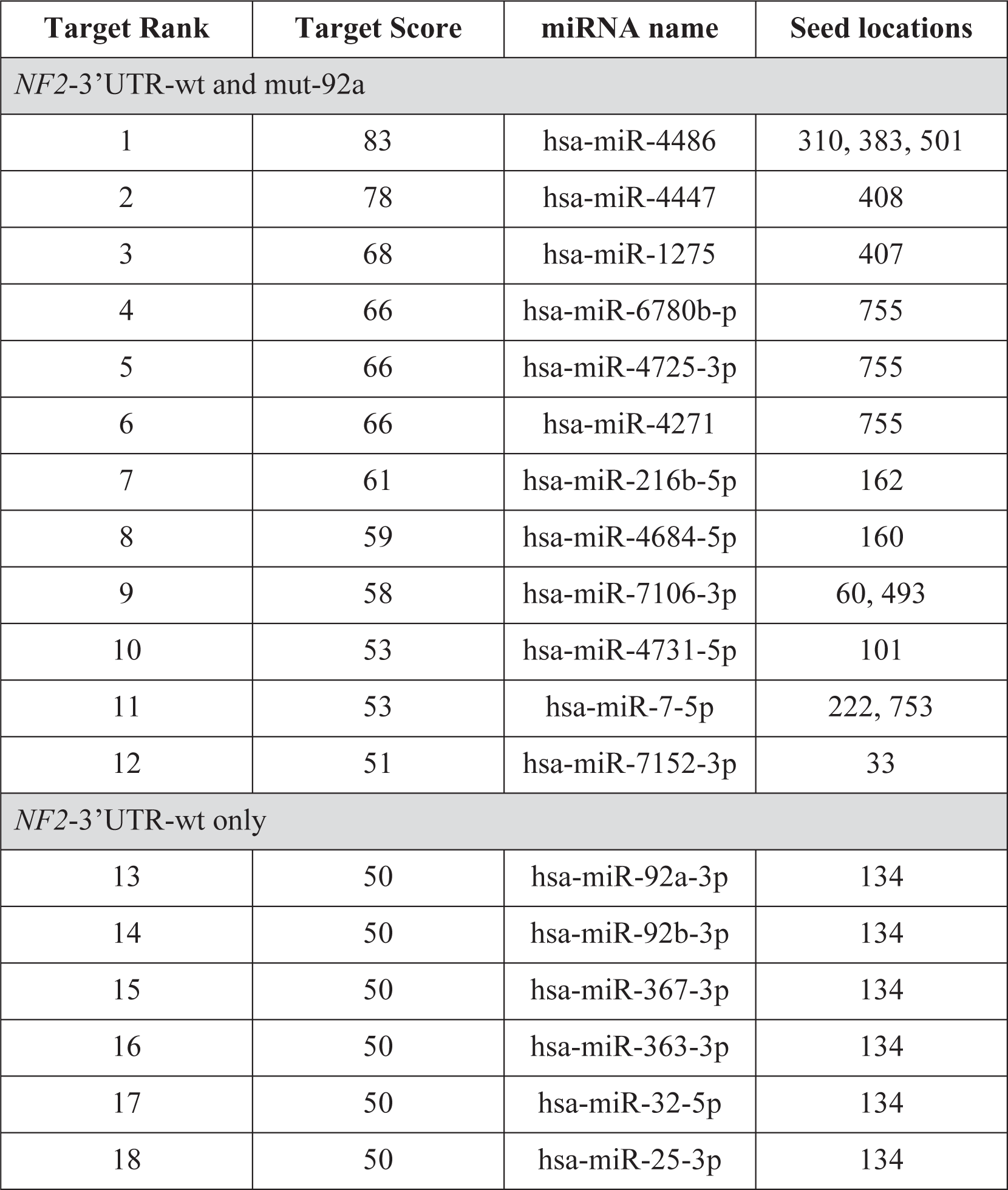
List of MirTarget predicted miRNAs that bind to the 792-bp cloned fragment of wild-type and miR-92a-MRE mutant *NF2*-3’UTR.

| **Target Rank** | **Target Score** | **miRNA name** | **Seed locations** |
| --- | --- | --- | --- |
| *NF2-3'UTR-wt and mut-92a* |  |  |  |
| 1 | 83 | hsa-miR-4486 | 310, 383, 501 |
| 2 | 78 | hsa-miR-4447 | 408 |
| 3 | 68 | hsa-miR-1275 | 407 |
| 4 | 66 | hsa-miR-6780b-p | 755 |
| 5 | 66 | hsa-miR-4725-3p | 755 |
| 6 | 66 | hsa-miR-4271 | 755 |
| 7 | 61 | hsa-miR-216b-5p | 162 |
| 8 | 59 | hsa-miR-4684-5p | 160 |
| 9 | 58 | hsa-miR-7106-3p | 60, 493 |
| 10 | 53 | hsa-miR-4731-5p | 101 |
| 11 | 53 | hsa-miR-7-5p | 222, 753 |
| 12 | 51 | hsa-miR-7152-3p | 33 |
| *NF2-3'UTR-wt only* |  |  |  |
| 13 | 50 | hsa-miR-92a-3p | 134 |
| 14 | 50 | hsa-miR-92b-3p | 134 |
| 15 | 50 | hsa-miR-367-3p | 134 |
| 16 | 50 | hsa-miR-363-3p | 134 |
| 17 | 50 | hsa-miR-32-5p | 134 |
| 18 | 50 | hsa-miR-25-3p | 134 |

**Table EV3.**
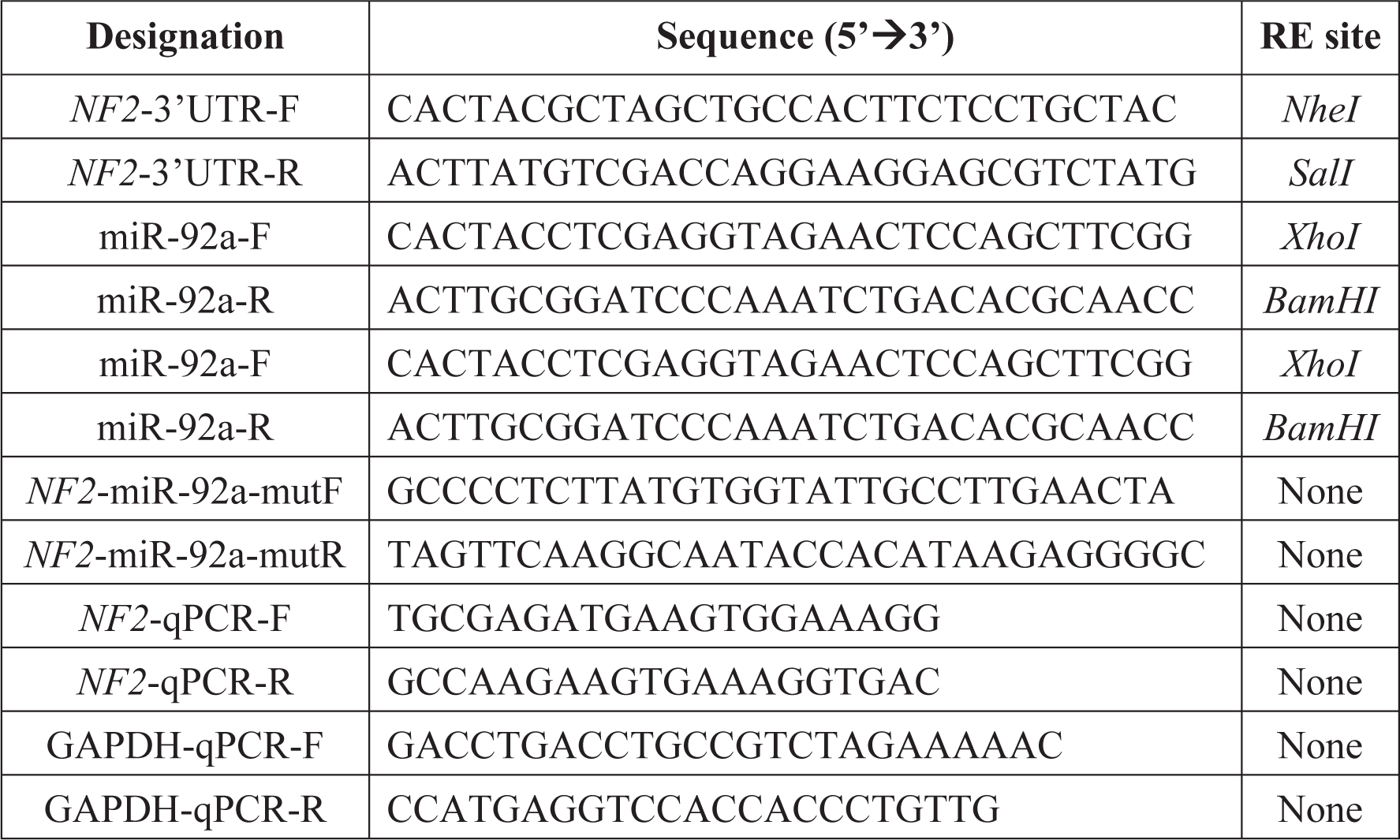
Primers used for construct generation, site-directed mutagenesis, and qPCR.

